# Scalable approximate Bayesian inference for particle tracking data

**DOI:** 10.1101/276253

**Authors:** Ruoxi Sun, Liam Paninski

## Abstract

Many important datasets in physics, chemistry, and biology consist of noisy sequences of images of multiple moving overlapping particles. In many cases, the observed particles are indistinguishable, leading to unavoidable uncertainty about nearby particles’ identities. Exact Bayesian inference is intractable in this setting, and previous approximate Bayesian methods scale poorly. Non-Bayesian approaches that output a single “best” estimate of the particle tracks (thus discarding important uncertainty information) are therefore dominant in practice. Here we propose a flexible and scalable amortized approach for Bayesian inference on this task. We introduce a novel neural network method to approximate the (intractable) filter-backward-sample-forward algorithm for Bayesian inference in this setting. By varying the simulated training data for the network, we can perform inference on a wide variety of data types. This approach is therefore highly flexible and improves on the state of the art in terms of accuracy; provides uncertainty estimates about the particle locations and identities; and has a test run-time that scales linearly as a function of the data length and number of particles, thus enabling Bayesian inference in arbitrarily large particle tracking datasets.

## 1. Introduction

In many biological and physical experiments it is necessary to track the movement of many isolated particles in a video datastream. This is an essential task in biomedical research, for example, to reveal the biophysical properties of both the imaged particles (e.g., single molecules) and the biological substrate (e.g., cell membrane) that the particles are traversing. Effective particle tracking algorithms have wide applications in both fundamental and applied biology, and more generally in chemistry and physical applications.

Previous scalable approaches to this task have largely involved non-Bayesian methods aiming at estimating a single “best” path of the underlying particles. However, in many applications particles have indistinguishable shapes under light microscopic resolution. This leads to a fundamen intal non-identifiability: if two particles pass close by each other (“meet”) then it is impossible to deterministically link the pre-meeting paths with the correct post-meeting paths (see Figure 1 below for an illustration). This motivates a Bayesian approach for assigning posterior probabilities over all the possible sets of particle paths consistent with the observed data.

**Figure 1.**
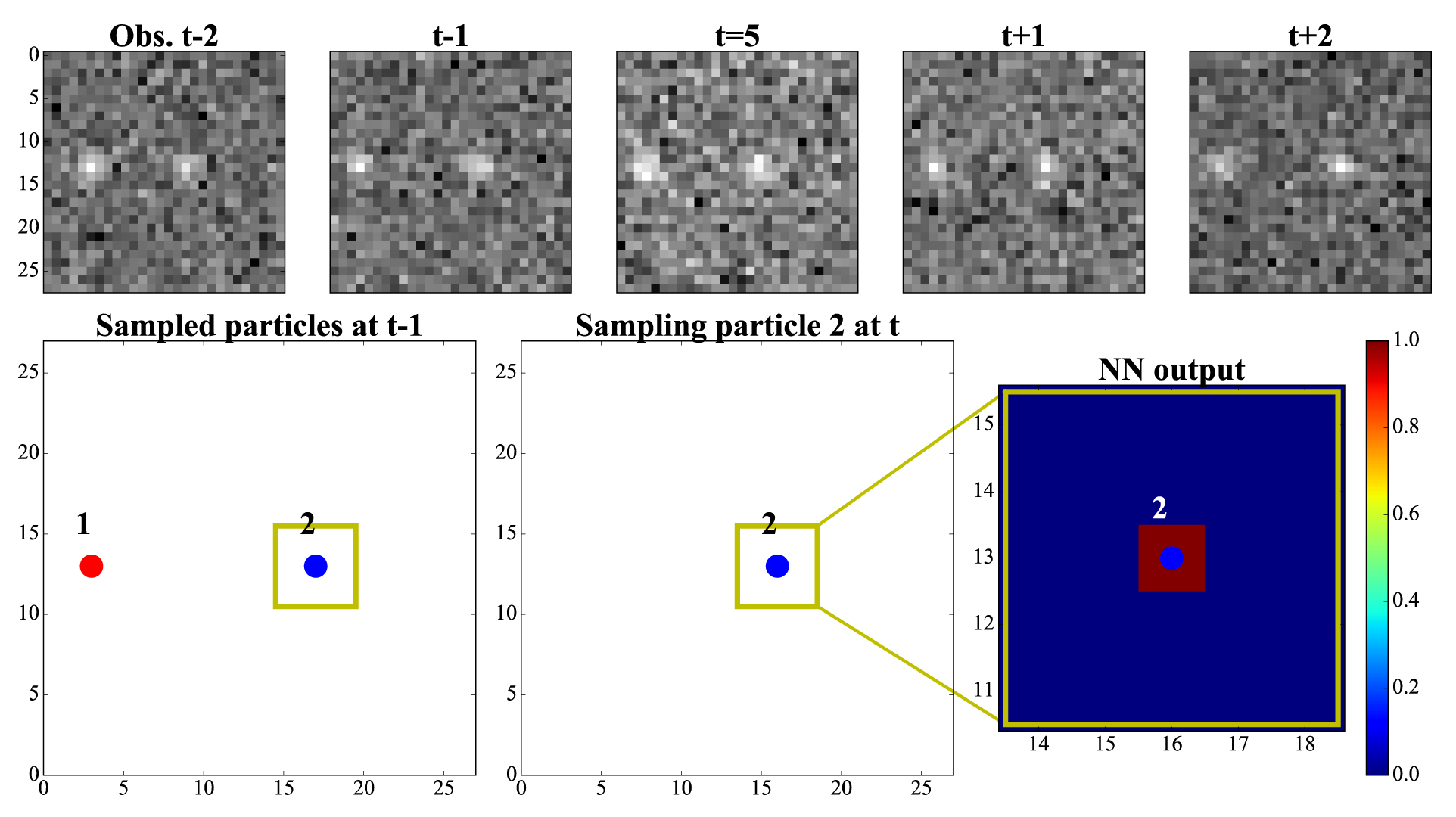
Overview of the conditional transition density network. Inputs to the network include the observed data *Y*_*t – M*:*t*+*M*_, with *M* = 2 here (top), the locations of particles sampled at time *t –* 1 (lower left), the particles that have been sampled so far at time *t* (lower middle), and the identity of the particle we are currently sampling (indicated by the yellow box). The network outputs the probability that the sampled particle survives to time *t*, and the conditional probability density of the particle’s next position. See the sampling process video for further illustration of the network processing data. In this video, the particles are restricted to move in the horizontal direction only (to facilitate plotting of the results in the following section); different particles are marked by different colors. The lower right panel displays the probability map 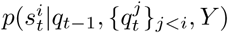 output by the network at each iteration.

Formally, at each timestep we observe a noisy, blurry image recording the particles’ current positions. In the simplest case, we can cast the tracking task in a factorial hidden Markov Model (**HMM**) framework, where each particle evolves according to a Markov process and thus multiple HMMs (one per particle) jointly determine the observed image data. The classic HMM inference approach is the forward-backward algorithm (Rabiner, 1990), but the complexity of forward-backward scales superlinearly with the number of particles here.

In this work, we propose an amortized inference approach utilizing a specialized recurrent neural network architecture to approximate the posterior particle transition densities inferred by forward-backward. After network training, posterior inference can be performed very quickly: given a new video dataset, the network outputs the conditional particle initialization and transition densities, and then we can simply sample forward from the resulting Markov chain to draw samples from the posterior particle paths.

We apply the method to simulated and real data. We show that the method robustly performs approximate Bayesian inference on the observed data, and provides more accurate results than competing methods that output just a single “best” path. Our approach is much more scalable than previously proposed Bayesian approaches, scaling linearly in the number of frames and in the number of observed pixels.

## 2. Model

We begin by describing the simplest concrete model for particle tracking data; we will generalize this below. We have *J* indistinguishable particles: each particle *j* appears at some time 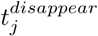 and disappears at some later time 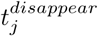 The particles move according to independent Gauss-Markov processes, with no interactions between particles. On each frame *t* we observe a blurred noisy sum of the particles that are visible at time *t*. The observation likelihood depends on the details of the experimental setup; the most common model is the Gaussian blur + Poisson noise model:

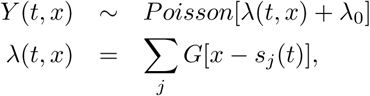

where *Y* (*t, x*) denotes the image data observed at pixel *x* at time *t, λ*_0_ is a background “dark noise” Poisson intensity, *G*[.] is a Gaussian point spread function (psf), *s*_*j*_(*t*) represents the location of particle *j* at time *t*, and the sum is over all particles that are alive at time *t*.

The model described above is a factorial HMM (Ghahramani & Jordan, 1996). However, this simple model can be generalized significantly. There may be multiple distinguishable classes of particles that have different shapes or colors. In many datasets particles can interact: they might merge, collide, split, etc. Individual particles often move in a non-Markovian manner (e.g., switching between several different latent dynamical modes). There may be strong dependencies between the motion of different particles, due e.g. to substrate motion. Finally, the observation noise may be highly non-Poisson, with correlations and strong inhomogeneities across the field of view. Thus it is critical to develop flexible inference approaches that do not depend on strong factorial HMM assumptions.

## 3. Related work on particle tracking

The literature on particle tracking methods is vast, and dates back to early physics studies of Brownian motion in fluids; see e.g. (Manzo & Garcia-Parajo, 2015) for a review, and (Chenouard et al., 2014) for a quantitative comparison of many algorithms. We will not attempt to review all of these methods here, but note that many algorithms split the tracking problem into a “detect” followed by a “link” step. The “detect” step outputs estimated particle locations given each image *Y*_*t*_. Various nonlinear filtering, thresholding, deconvolution, and neural network approaches have been employed for this task (Chenouard et al., 2014). Most such detection algorithms take just single frames *Y*_*t*_ as input, and therefore they do not integrate useful information across multiple frames to perform detection; (Newby et al., 2017) is a recent counterexample that demonstrates that better performance can be achieved if multiple frames *Y*_*t*_ are utilized in the detection step.

The “link” step then attempts to fuse these detected locations, to estimate the tracks that each visible particle took over the length of the observed movie. This linkage step is solved by some matching algorithm; see e.g. (Jaqaman et al., 2008) for an influential example of this approach, and (Chenouard et al., 2014; Wilson et al., 2016) for discussion of other linking methods.

As we emphasized in the introduction, deterministic detection and linking approaches are statistically suboptimal, since they ignore the irreducible uncertainty of the tracking problem that results when two or more visibly indistinguishable particles pass closer than a fraction of a psf-width of each other. Ignoring this uncertainty leads to non-robust results, in which tiny changes to the data can lead to discontinuous changes in the estimated particle tracks. Moreover, it is clear that the linkage and detection should not be separated: if we know the tracks of particles at times (1: *t –* 1) and (*t* + 1: *T*), then we have very strong prior information about the locations of particles at time *t*, and ignoring this useful prior information will lead to suboptimal results. (See e.g. (Sun et al., 2017), where similar points were made in the context of a related image-processing application.)

Similar points have been made in the Bayesian signal processing literature; for example, sequential Monte Carlo (particle filtering) methods have been applied to perform probabilistic inference in this setting (Smal et al., 2008). These approaches have the advantage of a proper grounding in standard Bayesian computational methodology, but scale poorly in the number of visible particles.

Finally, there is also a very large literature on “multi-target tracking,” e.g., tracking multiple people visible on security cameras. In this literature the different targets are typically distinguishable (e.g., different people visible on a camera will have different faces, gaits, clothing, etc.), whereas in this paper we focus on the case that the particles to be tracked are indistinguishable. Of course a middle ground exists in which particles have some distinguishing features but some posterior uncertainty about particle identity remains due to noisy or incomplete observations; however, to keep our presentation simple we focus exclusively on the most challenging fully-indistinguishable case here.

## 4. Methods

### 4.1 Overview

Our conceptual starting point is the standard filter-backward-sample-forward algorithm for sampling from the posterior distribution *p*(*Q |Y*) of the hidden state *Q* = {*q*_*t*_} of an HMM conditional on the observed data *Y* (Rabiner, 1990). This algorithm has two steps: (1) combine the observed data *Y* with the prior distribution *p*(*Q*) of the hidden Markov state *Q* to obtain a new Markov chain *p*(*Q*/*Y*), and (2) sample forward from this new Markov chain. Once (1) is complete, we can call (2) as often as we like to generate new sample paths from *p*(*Q|Y*).

This approach is attractive in our setting because sampling forward from a Markov chain is a fast operation once the conditional initial and transition densities (*p*(*q*_1_*|Y*) and *p*(*q*_*t*_*|Y, q*_*t-*1_), respectively) are in hand, where the hidden state *q*_*t*_ is the configuration of the locations and identities of all of the particles alive at time *t*. Thus in principle we can simply run (2) repeatedly to compute probabilities of any quantity we care about (e.g., the probability that a particle is in location *x* at time *t*, or the probability that particle *i* in frame *s* should be linked with particle *j* in frame *t*).

Unfortunately, as emphasized above, computing (1) exactly is intractable in our context; thus we need to approximate the conditional initial and transition densities. Our strategy is to train neural networks to approximate these probabilities. This approach is highly flexible; given enough training data, we can handle a wide variety of non-standard data, well beyond the simplest Gaussian blur + Poisson noise factorial HMM described above, since the learned probabilities do not lean heavily on special assumptions about e.g. the noise model or the precise details of the graphical model underlying the data^1^ In turn, we can generate as much training data as we need by simulating ground truth particle tracks along with the resulting observed data videos *Y*.

It is convenient to split the network into three parts: the conditional transition density that governs how samples move from timestep *t* to *t* + 1; the conditional birth density that governs the probability that a new particle appears at time *t*; and the conditional initial density that governs the positions of the particles at timestep 1. We describe each of these in turn below.

### 4.2 Conditional transition density network

This network is illustrated in Figure 1. The task of this network is to combine the observed data *Y* with the previous particle configuration *q* _*t-*1_ and to output probabilities that govern the. particle configuration *q*_*t*_ in the next time step This is a nontrivial task, since the dimensionality of *q*_*t*_ can be large and varies with time *t* as particles appear or disappear. Similarly, the observed image *Y*_*t*_ is often large (hundreds of pixels on a side), and in principle we need to observe multiple frames before and after time *t* to perform optimal inference.

Thus, for scalability, we break the problem up into a sequence of smaller pieces and work convolutionally. We begin by choosing a random ordering of the particles in *q*_*t-*1_. Then, for each of these particles indexed by *i*, we input three types of data: (1) a local patch of the observed movie data (in a spatial neighborhood around the *i*-th particle location 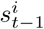, and in a temporal context of *M* frames before and after the current frame *t*; (2) a binary mask indicating the locations of the particles at time *t -* 1 in the same spatial neighborhood as particle *i*; and (3) a second binary mask indicating the locations of the particles *j* that have been sampled at time *t* prior to sampling particle *i*^2^. The network is then trained to output a probability map 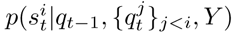 indicating the likely location 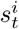 along with an auxiliary probability that the particle disappears (and is therefore no longer present at time *t*). Once these transition probabilities are learned, we can sample forward one particle and time-step at a time, as illustrated in the sampling process video and detailed in Algorithm 1; thus at test time inference scales linearly in the number of particles and time steps in the movie.

Note that we have slightly diverged from the vanilla filter-backward-sample-forward algorithm, which propagates information all the way back from the final observation *Y*_*T*_ to determine the state *q*_*t*_. Instead, we exploit the fact that only a local context around time *t* is necessary to infer *q*_*t*_, and thus we restrict our attention at time *t* to the local context *Y*_*t-M*:*t*+*M*_. (We use *M* = 2 throughout this paper.)

### 4.3 New birth networks and initialization

The network described above moves particles forward from timestep *t-*1 to *t*, and decides which particles should disappear at time *t*. However, new particles can enter the field of view at any time, and therefore we need a method for adding new particles to *q*_*t*_. Thus after running the update described above to *q*_*t*_, we run a second convolutional network that takes the same inputs as above (i.e., the local context of *q*_*t-*1_, *q*_*t*_, and *Y*, now at each location in the image instead of just at the previously-sampled particle locations) and outputs the probability that a new particle is born at each location at time *t*. Then we iteratively sample from this density and update *q*_*t*_ until no further particles are added.

#### Algorithm 1 Conditional sampling network

~~~
**Initialize:** *S*_1_ = Initializer(*Y*_1:*M*+1_, []) # **Sec. 4.3**
**# Sec. 4.2**
**for** *t* = 2; 3; 4::: **do**
      *S*_t_ = []
      **for** *i* in Permutation{*S*_t–1_} **do**
          *p*_*i*_ = ConditionalProbability(*Y*_*t*.*M*:*t*+*M*_, *S*_*t*–1_; *S*_t_, *i*)
          particle *i* disappears with prob. 1 – ∫ *p*_*i*_
          otherwise *i*′ is sampled from *p*_*i*_
          Insert *i*′ to *S*_*t*_
      **end for**
      *N*_*t*_ = NewBirth(*Y*_*t*..*M*:*t*+*M*_, *S*_*t*..1_; *S*_t_) # **Sec. 4.3**
      Insert *N*_*t*_ to *S*_*t*_
      *S*_t–1_ = *S*_t_
**end for**
~~~

The same strategy can be used to initialize *q*_1_; the only difference is that the inputs now don’t include *q*_*t-*1_ or the context of *Y* prior to *Y*_1_.

### 4.4 Network architecture and training

To handle the temporal and spatial dependencies in this data, we chose a combination of bi-directional 2D convolution LSTM and 3D convolutional layers; see Appendix. Overall, when the network is sampling forward, we can think of the resulting algorithm as a recurrent neural network (since the sampled output is then read back into the network to define the next state transition), with the somewhat non-standard feature that the network remains at timestep *t* for a random number of iterations (depending on how many particles need to be updated and how many particles are born at each timestep).

To train the network we generated simulated ground truth particle tracks *q*_*t*_ and corresponding observed movies *Y*. (We will discuss the training data in more detail in the following section.) Then we formed minibatches of training data, where each data sample included the inputs to the network (the local context of *Y*, *q*_*t-*1_, and a random subset of *q*_*t*_) along with the true particle location *s*^*i*^ (*t*), which served as the target output of the network. We trained the network (using default learning rate settings in Keras) to minimize the binary cross-entropy between the target mask (zero except at *s*^*i*^(*t*), or all zeros if all the particles in *q*_*t*_ were already sampled and no further particles should be added) and the network’s output probability mask.

## 5. Results

### 5.1 One-dimensional example

We begin with a simple simulated experiment in which the particles are restricted to move in the horizontal direction only. This makes it easier to view and understand the results, by simply plotting the horizontal positions of the (true vs. inferred) particles as a function of time. The results are illustrated in Figure 2; the same data are shown in Figure 1 and the sampling process video.

**Figure 2.**
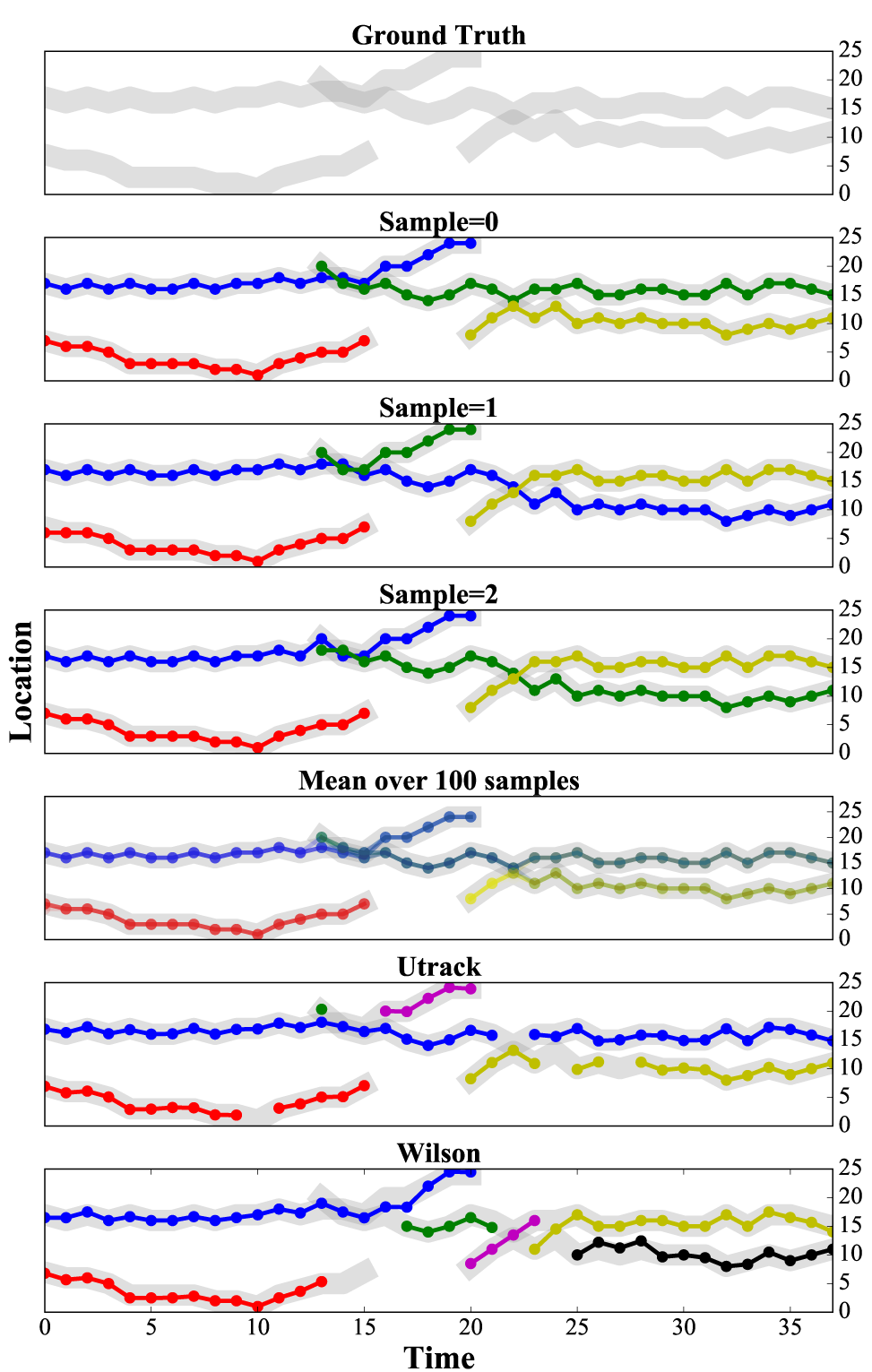
One-dimensional simulated example. We test the performance of the proposed algorithm on a simplified example where the particles are restrained to move in one dimension, to facilitate visualization. See the sampling process video for the raw data. **Top**: Ground truth tracks. New particles appear near *t* = 13 and *t* = 20; a particle disappears near *t* = 20; “meetings” between two particles occur near *t* = 14 and *t* = 22. **Panels 2-4**: sample tracks output by our proposed method. Colors indicate particle identity. Note that the detected locations track the ground truth locations and appearance/disappearance times accurately, and identity is assigned probabilistically following particle meetings, as desired. (The network output corresponding to Sample 0 is shown in Figure 1 and the sampling process video, with colors matched across the figures and video.) **Panel 5**: mean over 100 examples; the blended colors following particle meeting times indicate the relative probabilities of the identity assignments. **Bottom two panels**: output from two deterministic particle tracking methods, by (Jaqaman et al., 2008) and (Wilson et al., 2016), respectively. Several detection errors are visible in the output of these methods, leading to oversegmentation of the output tracks.

In this example we see the appearance and disappearance of a couple particles, and two “meeting” events in which one particle overlaps significantly with another particle. Since in this example all the particles have identical shapes and are undergoing independent and identically distributed Brownian motions, there is no way to deterministically “link” particles before and after these meeting events; i.e., the “correct” linker here must output a probabilistic answer.

In panels 2-4 of Figure 2 we display three conditional sample paths drawn by our algorithm. Sample 0 (panel 2) recovers the ground truth accurately, and Sample 1 and 2 (panels 3 and 4) give different — but also valid — sets of tracks. Panel 5 shows an average of 100 samples overlaid together, with the colors indicating relative probabilities of the chosen tracks. Note that at the beginning of the trial, where the two visible particles are well-isolated, the sampler essentially outputs a deterministic estimate, with all samples assigned to the left (red) or the right (blue). However, after the “meeting” near *t* = 15, the colors blend, indicating probabilistic assignment of tracks following this event, as desired.

For comparison, we also show the output of two existing particle tracking methods, both of which output deterministic particle identities. Our approach provides visibly more robust outputs on this example, with fewer dropped particle detections and false particle appearances or disappearances.

### 5.2 Two-dimensional example

Next we turn to a small-scale simulated two-dimensional example; the results are illustrated in the moving particles video, Figure 3, and the 3D view video. As in the previous one-dimensional example, we find that our proposed approach accurately detects the particle locations and appearance/disappearance times, and successfully assigns identities probabilistically following particle meetings.

**Figure 3.**
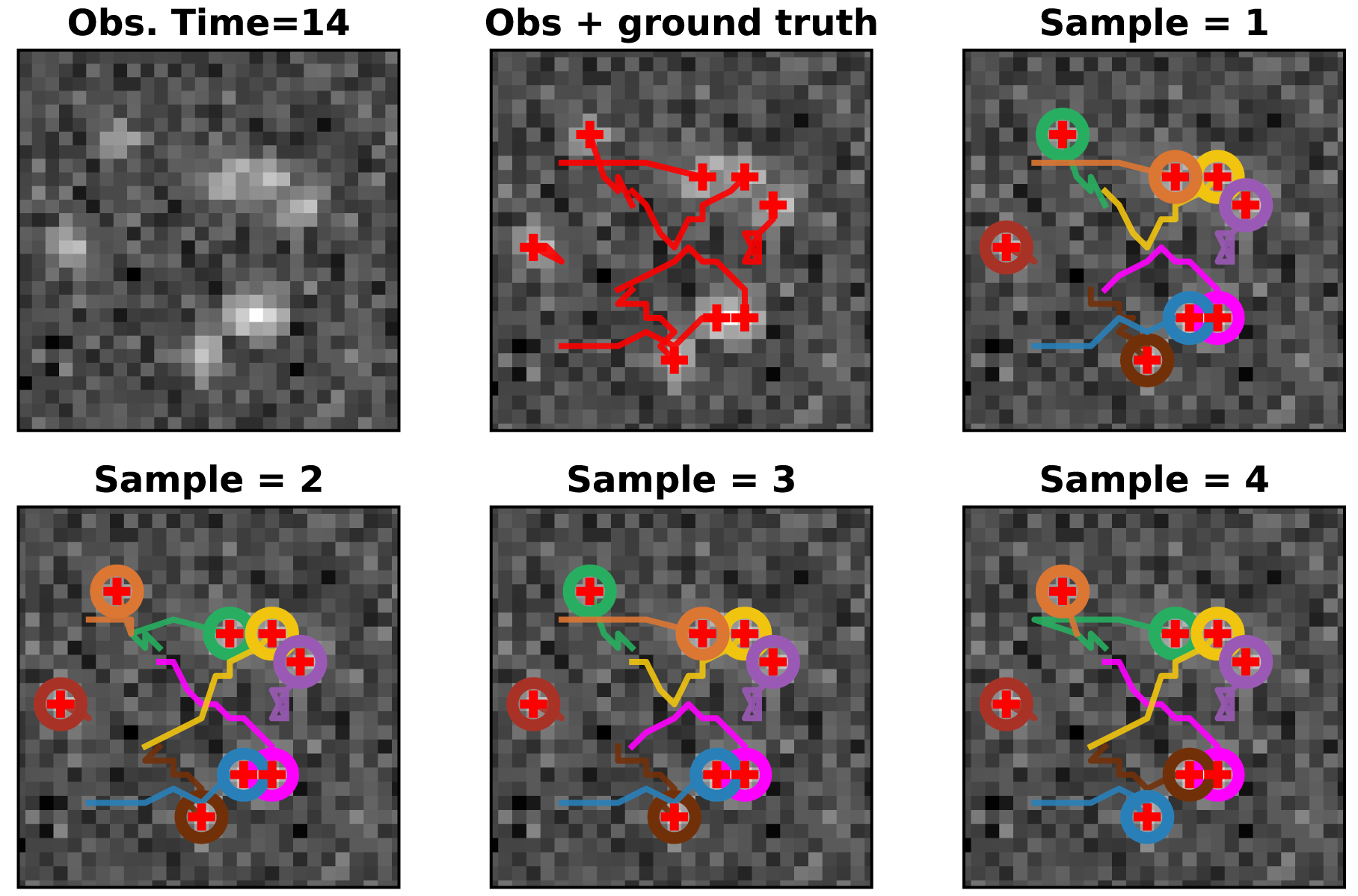
Two dimensional example. This figure displays a single frame of the moving particles video; view the video for further details. **First panel**: observed data *Y*_*t*_. **Top middle**: a time-lapse trace of the ground truth particle tracks; plus marks the current particle position, and the tails mark the recent history. **Other panels**: four sample estimated particle tracks output by our proposed method. As can be seen in the moving particles video, as particles meet the particle identities are assigned probabilistically and the identities across different samples diverge, even if some identities were initially the same across some samples. (Colors of sampled tracks are assigned according to total proximity of each track to the ground truth tracks, each of which is assigned a random color.) We show another representation of these samples in the 3D view video; the first frame of this video shows the ground truth tracks and each remaining frame shows a single sample in 3d (two spatial dimensions and one time dimension), with colors matched to those shown in this figure and in the moving particles video; thick lines indicate ground truth tracks for comparison.

### 5.3 Large scale examples and evaluation

To establish a more quantitative evaluation, we compared against two baseline methods: the popular Utrack approach (Jaqaman et al., 2008) and the method proposed in (Wilson et al., 2016), which performed well on the performance metrics established in the review / competition paper (Chenouard et al., 2014). We generated large-scale two-dimensional simulated data whose parameters matched a pair of challenging datasets in (Chenouard et al., 2014), and then computed the suite of performance metrics (measuring various facets of detection accuracy, linking quality, etc.) introduced in the same paper (averaging over 100 draws from our sampler for each dataset). Results are shown in Table 1: we find that our proposed method outperforms the baselines on both datasets examined, on all the performance metrics computed here.

**Table 1.** Comparison of three particle tracking methods: our proposed approach (“network”), Utrack from (Jaqaman et al., 2008), and the method proposed in (Wilson et al., 2016). Bold indicates best performance; we find that the proposed network approach achieves the best performance over both datasets and all performance metrics computed here. RMSE: Root Mean Square Error; COV: Coverage; JSC: Jaccard similarity coefficient; all quantities are as defined in (Chenouard et al., 2014).

|  | SNR=5 |  |  |  |  |  |  |
| --- | --- | --- | --- | --- | --- | --- | --- |
|  | Assign error | RMSE | COV | Alpha | Beta | JSC | JSC<sub>θ</sub> |
| Network | **7.28 ± 1.05** | **0.08 ± 0.01** | **0.69 ± 0.05** | **0.82 ± 0.06** | **0.81 ± 0.03** | **0.68 ± 0.06** | **0.99 ± 0.02** |
| Utrack | 8.88 | 0.189 | 0.574 | 0.778 | 0.748 | 0.523 | 0.769 |
| Wilson | 11.83 | 0.214 | 0.385 | 0.704 | 0.667 | 0.339 | 0.741 |
|  | SNR=4 |  |  |  |  |  |  |
|  | Assign error | RMSE | COV | Alpha | Beta | JSC | JSC<sub>θ</sub> |
| Network | **8.23 ± 1.12** | **0.07 ± 0.01** | **0.63 ± 0.07** | **0.76 ± 0.03** | **0.70 ± 0.04** | **0.55 ± 0.08** | **0.82 ± 0.05** |
| Utrack | 12.78 | 0.10 | 0.32 | 0.62 | 0.59 | 0.29 | 0.68 |
| Wilson | 14.42 | 0.34 | 0.36 | 0.58 | 0.55 | 0.32 | 0.63 |

It is worth emphasizing that these performance metrics were designed for deterministic tracking algorithms, and therefore entirely miss one of the major advantages of our approach (the fact that it outputs not just a single “best” track estimate but instead estimates the posterior distribution over all tracks). How can we evaluate the quality of our approximation to the posterior here (and quantitatively compare between different algorithms that attempt to approximate this posterior)? One natural approach is to estimate the Kullback-Leibler divergence *D*_*KL*_[*f* (*Q*); *p*(*Q Y*)] between our approximate posterior *f* (*Q*) and the true posterior *p*(*Q Y*) on the state space *Q* given the observed data *Y*. Of course, this is not quite tractable, due to the intractability of *p*(*Q Y*), but we can estimate *D*_*KL*_[*f* (*Q*); *p*(*Q Y*)] up to a constant in *f* (*Q*) by sampling from *f* (*Q*):

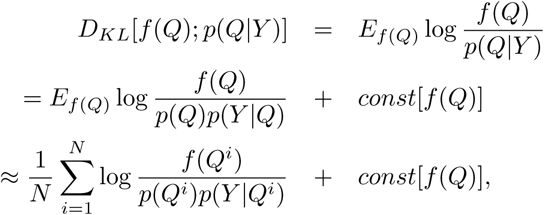

where *Q*^*i*^) _*i*=1:*N*_ are *N* samples from *f* (*Q*). Here *p*(*Q*^*i*^) and *p*(*Y Q*^*i*^) can be evaluated explicitly if the prior *p*(*Q*) is e.g. Markovian; for our approach *f* (*Q*^*i*^) can also be evalu ated directly since *f* (*Q*) has an explicit Markov form. This provides us a method for scoring any Bayesian particle tracking algorithm for which we can explicitly evaluate the approximate posterior *f* (*Q*). (We do not perform this scoring on the baselines examined here, since for any deterministic algorithm *f* (*Q*) is a delta function, leading to an infinite Kullback-Leibler score if we treat *q*_*t*_ as a continuous random variable — i.e., the probabilistic approach trivially outperforms deterministic approaches.)

### 5.4 Real data example

Finally, we tested the performance of our algorithm on real data. The data are TIR-FM imaged clathrin-coated pits in a BSC1 cell (Jaqaman et al., 2008). We trained the network on simulated data whose parameters (signal-to-noise ratio, particle density and speed, psf width, etc.) were coarsely matched to the real data; see the comparison video for details. We plot three samples from our algorithm using different colors in Fig. 4 and the real data video. While ground truth is unavailable in this case, by visual inspection the algorithm seems to effectively follow the particles in the video, without excessive oversegmentation of the tracks; the output here seems consistent with the behavior of the algorithm on the previous simulated datasets.

**Figure 4.**
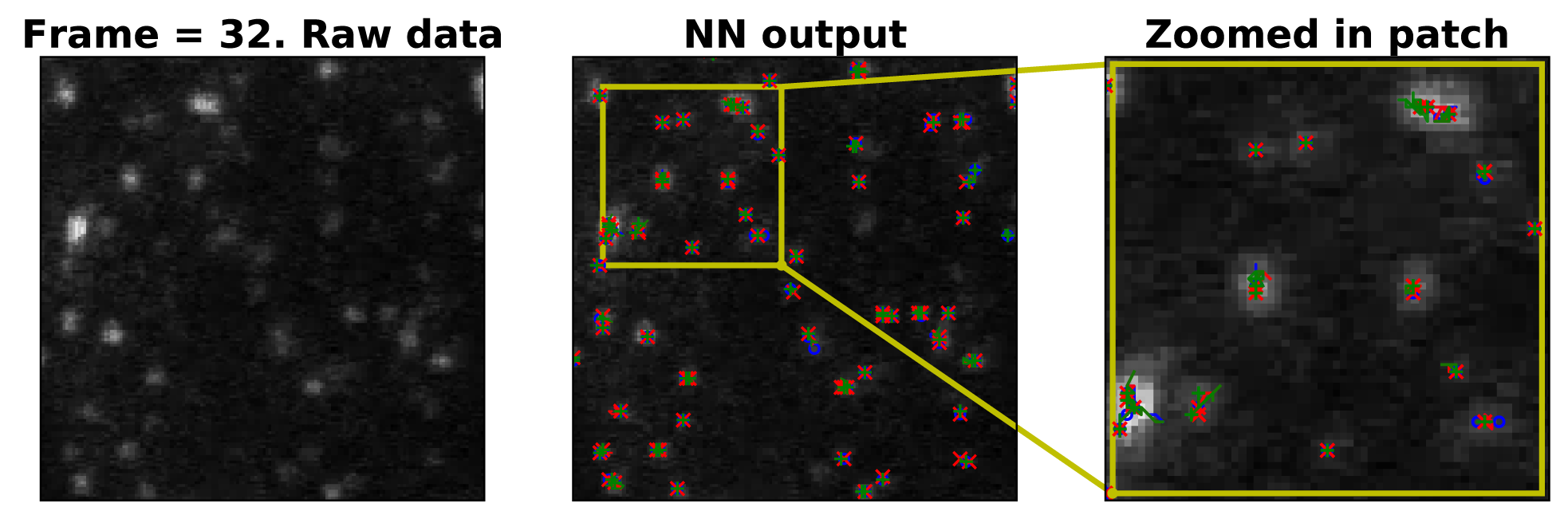
Real data. Performance on real data (TIR-FM imaged clathrin-coated pits in a BSC1 cell) (Jaqaman et al., 2008). **Left**: raw image sequence. **Middle**: raw image sequence overlaid with detection markers and tails indicating the recent location history. Colors indicate three different samples from our algorithm. **Right**: a zoomed in patch. Image size: 150 170 pixels, pixel size: 67 nm. For more details see the real data video.

**Figure 5.**
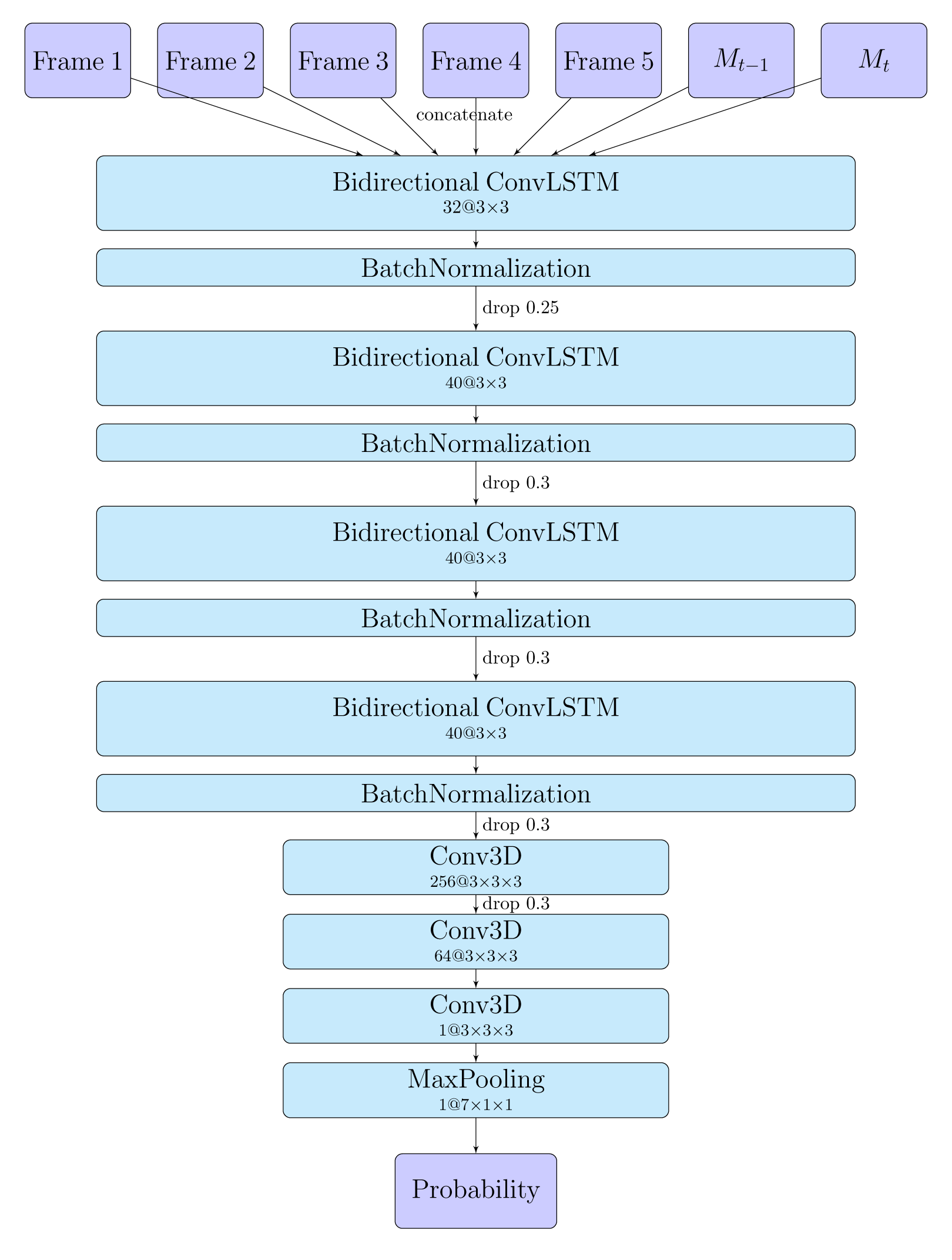
Neural network architecture. This is the architecture of the conditional transition density network described in section 4.2. We use Bidirectional Convolution LSTM and Convolution 3D Neural Networks as building blocks with notation ‘40@3×3’ meaning 40 feature maps with kernel size 3 by 3. The input size of the networks is 7 *× patch size × patch size*: 5 patches for *Y*_*t-*2:*t*+2_, plus binary masks *M*_*t-*1_ and *M*_*t*_ encoding the positions of the particles at times *t -* 1 and *t* respectively. The output probability map is *patch size × patch size*. We used *patch size* = 28 here. The new birth network in section 4.3 uses a similar architecture.

## 6. Discussion

### 6.1 Related machine learning work

In the introduction we emphasized the importance of the particle tracking problem; we believe that the more robust, accurate, and probabilistic tracking methods developed here will have a significant impact in a wide range of biological and physical applications.

More generally, from a machine learning point of view, the major novelty of our work is the incorporation of neural network methods to provide a flexible and scalable approximation of Bayesian inference via efficient sampling in a large graphical model.

Of course, interactions between Bayesian analysis and neural network methods comprise a very rich thread of research these days. The work of (Snell & Zemel, 2017) is highly relevant: this paper describes a neural network approach to sample multiple segmentations that are consistent with an observed image, much as we use neural networks to sample multiple particle tracks that are consistent with an observed video.

As another example, variational autoencoders (Kingma & Welling, 2013; Rezende et al., 2014) and variants thereof (Johnson et al., 2016; Gao et al., 2016; Fraccaro et al., 2017; Krishnan et al., 2017) have become very popular recently for performing inference in nonlinear HMMs. These methods are most effective when the latent state variable is lowdimensional. In the particle tracking problem the latent dynamical variable is very high-d (scaling with the number of particles) and more importantly the latent dimensionality is time-varying, as particles are born, die, merge, split, enter, or leave the focal plane. We are not aware of variational autoencoder approaches that would be easily applicable to the particle tracking problem.

Another related thread involves amortized inference using neural networks for sequential Monte Carlo (“particle filtering,” which is not to be confused with the particle tracking problems considered in our paper); see e.g. (Paige & Wood, 2016). Again, it is not clear how well these methods would scale to the large-scale multiple-particle tracking problems of interest here.

Finally, our work is an example of a broad theme in the current image processing literature: start with “ground truth” images, then simulate observed data that can be generated as some kind of corruption of this ground truth, and then use this simulated data to train a neural network that can “denoise” (or super-resolve, or deblur, or infill, etc.) this corruption. A (highly non-exhaustive) list of recent examples includes: (Parthasarathy et al., 2017), which applies this idea to approximate Bayesian decoding of neuronal spike train data; (Yoon et al., 2017), to segmentation of threedimensional neuronal images; and (Weigert et al., 2017), to denoising of microscopy images.

### 6.2 Future work

At test time, as emphasized above, the inference approach proposed here is highly scalable, but the network training time is relatively slow (taking on the order of hours for the experiments presented here). This is typical of “amortized inference” approaches: we pay with relatively long training times for fast test times. Thus our proposed approach is most valuable in settings where we have repeated experimental samples from a similar data regime (instead of training a new inference network for each new experimental dataset).

We have not expended serious effort optimizing over network architectures here; we could likely find lighter architectures that perform similarly, which would speed up both testing and training. Similarly, we could distill/compress the network to further speed up test times, if necessary e.g. for online experimental designs.

Similarly, we have not yet attempted to develop automated procedures for choosing parameters for generating training data. In practice we have found that these parameters (e.g., the amplitude, density, variance/speed of particles, plus noise levels, point-spread width, etc.) are fairly straightforward to choose, and the inference results are not highly sensitive to small misspecifications of these parameters (recall Figure 4 and the corresponding comparison video). It would be useful to develop a simple interface that would allow experimentalists to easily generate training data, followed by generation of a network trained to perform inference on their data.

An alternative approach would be to include data parameters as extra inputs for the network. Then in principle there would be no need to train a new network for each new type of data; instead we could perhaps just train a single big network on many different data types (with the corresponding data parameters included as inputs to the network) and then when presented with a new data type we just provide the network with the required parameters and let it perform inference. This is an ambitious but important direction for future work^3^.

## Footnotes

1 The main assumption we make is that the posterior *p*(Q|*Y*) can be well-approximated as Markovian, so that our resulting Markovian sampler can provide good approximations to true samples from the posterior. This assumption is reasonable in the majority of particle-tracking applications we have in mind.

2 This input lets the network avoid placing two particles to explain a single observed bump in *Y*_*t*_; if a previously-sampled particle *j* already explains the bump well, then the network will prefer to put particle *i* elsewhere. Also note that the input data *Y* and output probability maps don’t need to have the same number of pixels (i.e., we could attempt to resolve the particle locations at higher spatial resolution than the observed data), but we have not pursued this direction in detail.

3 Note that a slightly different philosophy is espoused in (Newby et al., 2017), who trained a single deterministic network for particle detection that can be applied to a wide range of data, but without including any parameters describing the data generation mechanism as inputs to the network. This approach makes it easy for experimentalists to use the network (since no training or parameter estimation is required), but likely sacrifices some accuracy compared to a network that is provided information about the parameters governing the generation of the data. We hope to run more detailed comparisons of these approaches in the future.

## References

Chenouard, Nicolas, Smal, Ihor, De Chaumont, Fabrice, Maška, Martin, Sbalzarini, Ivo F, Gong, Yuanhao, Cardinale, Janick, Carthel, Craig, Coraluppi, Stefano, Winter, Mark, et al. Objective comparison of particle tracking methods. Nature methods, 11(3):281, 2014.

Fraccaro, Marco, Kamronn, Simon, Paquet, Ulrich, and Winther, Ole. A disentangled recognition and nonlinear dynamics model for unsupervised learning. In Advances in Neural Information Processing Systems, pp. 3604–3613, 2017.

Gao, Yuanjun, Archer, Evan W, Paninski, Liam, and Cunningham, John P. Linear dynamical neural population models through nonlinear embeddings. In Lee, D. D., Sugiyama, M., Luxburg, U. V., Guyon, I., and Garnett, R. (eds.), Advances in lNeural Information Processing Systems 29, pp. 163–171. 2016.

Ghahramani, Zoubin and Jordan, Michael I. Factorial hidden markov models. In Advances in Neural Information Processing Systems, pp. 472–478, 1996.

Jaqaman, Khuloud, Loerke, Dinah, Mettlen, Marcel, Kuwata, Hirotaka, Grinstein, Sergio, Schmid, Sandra L, and Danuser, Gaudenz. Robust single-particle tracking in live-cell time-lapse sequences. Nature methods, 5(8): 695, 2008.

Johnson, Matthew, Duvenaud, David K, Wiltschko, Alex, Adams, Ryan P, and Datta, Sandeep R. Composing graphical models with neural networks for structured representations and fast inference. In Advances in neural information processing systems, pp. 2946–2954, 2016.

Kingma, Diederik P and Welling, Max. Auto-encoding variational bayes. arXiv preprint arXiv:1312.6114, 2013.

Krishnan, Rahul G, Shalit, Uri, and Sontag, David. Structured inference networks for nonlinear state space models. In AAAI, pp. 2101–2109, 2017.

Manzo, Carlo and Garcia-Parajo, Maria F. A review of progress in single particle tracking: from methods to biophysical insights. Reports on progress in physics, 78 (12):124601, 2015.

Newby, Jay M, Schaefer, Alison M, Lee, Phoebe T, Forest, M Gregory, and Lai, Samuel K. Deep neural networks automate detection for tracking of submicron scale particles in 2d and 3d. arXiv preprint arXiv:1704.03009, 2017.

Paige, Brooks and Wood, Frank. Inference networks for sequential monte carlo in graphical models. In International Conference on Machine Learning, pp. 3040–3049, 2016.

Parthasarathy, Nikhil, Batty, Eleanor, Falcon, William, Rutten, Thomas, Rajpal, Mohit, Chichilnisky, EJ, and Paninski, Liam. Neural networks for efficient bayesian decoding of natural images from retinal neurons. In Advances in Neural Information Processing Systems, pp. 6437–6448, 2017.

Rabiner, Lawrence R. Readings in speech recognition. chapter A Tutorial on Hidden Markov Models and Selected Applications in Speech Recognition, pp. 267–296. Morgan Kaufmann Publishers Inc., 1990.

Rezende, Danilo Jimenez, Mohamed, Shakir, and Wierstra, Daan. Stochastic backpropagation and approximate inference in deep generative models. arXiv preprint arXiv:1401.4082, 2014.

Smal, Ihor, Draegestein, Katharina, Galjart, Niels, Niessen, Wiro, and Meijering, Erik. Particle filtering for multiple object tracking in dynamic fluorescence microscopy images: Application to microtubule growth analysis. IEEE Transactions on Medical Imaging, 27(6):789–804, 2008.

Snell, Jake and Zemel, Richard S. Stochastic segmentation trees for multiple ground truths. In Proceedings of the Thirty-Third Conference on Uncertainty in Artificial Intelligence, 2017.

Sun, Ruoxi, Archer, Evan, and Paninski, Liam. Scalable variational inference for super resolution microscopy. In Singh, Aarti and Zhu, Jerry (eds.), Proceedings of the 20th International Conference on Artificial Intelligence and Statistics, volume 54 of Proceedings of Machine Learning Research, pp. 1057–1065. PMLR, 2017.

Weigert, Martin, Schmidt, Uwe, Boothe, Tobias, Andreas, M, Dibrov, Alexander, Jain, Akanksha, Wilhelm, Benjamin, Schmidt, Deborah, Broaddus, Coleman, Culley, Siân, et al. Content-aware image restoration: Pushing the limits of fluorescence microscopy. bioRxiv, pp. 236463, 2017.

Wilson, Rhodri S, Yang, Lei, Dun, Alison, Smyth, Annya M, Duncan, Rory R, Rickman, Colin, and Lu, Weiping. Automated single particle detection and tracking for large microscopy datasets. Royal Society open science, 3(5): 160225, 2016.

Yoon, Young-Gyu, Dai, Peilun, Wohlwend, Jeremy, Chang, Jae-Byum, Marblestone, Adam H, and Boyden, Edward S. Feasibility of 3d reconstruction of neural morphology using expansion microscopy and barcode-guided agglomeration. Frontiers in computational neuroscience, 11, 2017.

